# Mechanistic insights and *in vivo* HIV suppression by the BRD4-targeting small molecule ZL0580

**DOI:** 10.1101/2025.08.14.670267

**Authors:** Naveen Kumar, Zonghui Ma, Fuquan Long, Srinivasa Reddy Bonam, Hsien-Tsung Lai, Shwu-Yuan Wu, Haiying Chen, Nicholas C. Hazell, Jiani Bei, Xuefeng Liu, Zhi Wei, Cheng-Ming Chiang, Jia Zhou, Haitao Hu

## Abstract

Epigenetic suppression and durable silencing of HIV represent a promising strategy to achieve ART-free remission, consistent with the “block and lock” HIV cure paradigm. BRD4 is a host epigenetic reader and plays a critical role in HIV transcriptional regulation. We previously identified ZL0580, a first-in-class BRD4-selective small molecule distinct from the pan-BET inhibitor JQ1, which induces HIV epigenetic suppression. However, detailed molecular mechanisms, pharmacokinetics (PK), and *in vivo* HIV-suppressive efficacy of ZL0580 remain undefined. Here, we show that ZL0580 selectively targets BRD4 bromodomain 1 (BD1) through interaction with a key glutamic acid residue (E151), as determined by structural modeling and mutagenesis. Transcriptomic profiling by RNA-seq reveals that ZL0580 and JQ1 induce opposing gene expression programs, consistent with their distinct effects on HIV proviral transcription and latency. In a humanized mouse model of HIV infection, ZL0580 monotherapy, or in combination with ART, potently suppressed active HIV replication, reducing the plasma viremia to nearly undetectable levels, and delayed viral rebound following treatment interruption. Collectively, these findings establish ZL0580 as an epigenetic suppressor of HIV *in vivo* and provide proof-of-concept for its potential as a “block and lock” HIV cure candidate, warranting further optimization and development.

## INTRODUCTION

Human immunodeficiency virus (HIV) persists in latent reservoirs that evade immune surveillance and remains refractory to antiretroviral therapy (ART), posing a major barrier to an HIV cure [1–3]. Although ART effectively suppresses active viral replication and reduces plasma viremia to undetectable levels [4], it does not eradicate latent reservoirs, which are the primary source of viral persistence [5–7]. Consequently, treatment interruption almost invariably results in rapid viral rebound [3, 8, 9] and intensification of ART is ineffective in reducing reservoir size or residual viral transcription [10]. These limitations highlight the need for novel therapeutic strategies aimed at achieving HIV remission or cure [5, 11].

HIV proviral transcription and latency are tightly controlled by host epigenetic and transcriptional mechanisms [12–14]. Unlike the “shock and kill” strategy, which seeks to reverse latency and eliminate infected cells [15–17], the “block-and-lock” approach aims to achieve a functional cure by reinforcing latency and preventing viral reactivation through modulation of host and viral pathways [18–20]. For examples, LEDGIN, an HIV integrase inhibitor, hampers viral transcription and prevents viral reactivation by retargeting the provirus to transcriptionally inactive sites [21–23]. The Tat inhibitor didehydro-cortistatin A (dCA) suppresses Tat-dependent transactivation and promotes deep latency by inducing epigenetic repression in the HIV promoter [24–26]. In HIV-infected humanized mouse models, dCA treatment reduced residual viral replication and delayed viral rebound following treatment interruption [27].

BRD4 is a host cell epigenetic reader belonging to the bromodomain (BD) and extra-terminal (ET) domain protein family (BET) [28, 29]. It is characterized by two conserved tandem bromodomains (BD1 and BD2) and an ET domain [30–33]. A primary function of BRD4 is to act as a scaffold by binding to acetylated histones and recruiting partner proteins to the gene promoters, thereby modulating gene expression [34]. The role of BRD4 in regulating HIV transcription and latency has been documented [30, 35–37]. Inhibition of BET/BRD4 by the pan-BET inhibitor JQ1 has been shown to activate HIV transcription and reverse latency [35–37]. However, evidence from the studies of our group and others suggests that BRD4 has versatile function and exerts context-dependent effects on HIV transcription, largely determined by its specific interactions with histones and partner proteins [38, 39].

In our previous studies, we identified ZL0580, a first-in-class BRD4-selective small molecule distinct from the pan-BET inhibitor JQ1, which induces HIV epigenetic suppression and promotes deeper state of latency in cellular models [38, 40, 41]. This effect has been demonstrated in multiple *in vitro* and *ex vivo* models, including J-Lat cells, primary CD4⁺ T cells, and peripheral blood mononuclear cells (PBMCs) from ART-suppressed individuals, as well as myeloid cells such as microglia and macrophages [38, 40, 41]. ZL0580 suppresses HIV by altering chromatin structure at the viral LTR and inhibiting Tat-mediated transcription elongation [38, 40, 41]. The HIV-suppressive effect of ZL0580 was recently confirmed by another study [23]. Despite these findings, the molecular basis of the selective interaction of ZL0580 and BRD4 (as compared with JQ1), its pharmacokinetics (PK), and *in vivo* HIV-suppressive activity (block-and-lock potential) remain undefined.

In this study, we addressed several gaps critical for advancing ZL0580 toward further development. First, molecular docking and protein mutagenesis identified the glutamic acid at position 151 in the BD1 of BRD4 as a key residue that mediates the selective ZL0580-BRD4 interaction. Second, RNA-seq analysis revealed that ZL0580 and JQ1 induce largely opposing transcriptomic profiles, consistent with their distinct effects on HIV transcription and latency. Third, we characterized *in vivo* PK and safety profiles of ZL0580 in rodents, which, together with its HIV-suppressive activity demonstrated in diverse *in vitro* and *ex vivo* models, support its therapeutic potential. Finally, in a humanized mouse model of HIV infection, we demonstrate that ZL0580 monotherapy, or its combination with ART, suppresses active HIV replication, reducing plasma viremia to nearly undetectable levels after two weeks of treatment, and also delays viral rebound after treatment interruption, providing early proof-of-concept evidence for its *in vivo* block-and-lock activity.

## RESULTS

### Distinct binding modes of ZL0580 *vs*. (+)-JQ1 to human BRD4 BD1 domain

ZL0580 was identified as a selective BRD4 BD1 inhibitor in our previous studies (**Fig. 1A**) [38]. To elucidate its molecular interactions with BRD4 BD1, we performed molecular docking study based on the co-crystal structure of BRD4 BD1 in complex with ZL0590, a close analog of ZL0580 [42]. The co-crystal structure of ZL0590 complexed with BRD4 BD1 was used for molecular simulation study to explore the key interactions between ZL0580 and BRD4 BD1 [42]. Docking results revealed that ZL0580 adopts a binding mode similar to ZL0590. As shown in **Fig. 1B-C**, ZL0580 forms several direct and water-mediated indirect hydrogen bond interactions with key residues in BRD4 BD1 protein, which are predicted to contribute to the high binding affinity and selectivity of ZL0580 for BRD4 BD1. Specifically, ZL0580 forms two water-mediated hydrogen bonds with the carbonyl group of GLU154 residue through two NH groups of the urea moiety. Moreover, the carbonyl oxygen of the urea moiety forms a water-mediated hydrogen bond with the hydroxyl group of TYR137. Two oxygen atoms of the sulfonyl group of ZL0580 form direct and water-mediated hydrogen bond interactions with the NH group of GLY143 residue, respectively. Additionally, the NH group of the amide moiety forms a direct hydrogen bond with the carbonyl oxygen atom of GLU151 residue (**Fig. 1B-C**). Unlike JQ1, ZL0580 targets a novel binding site spatially distinct from the classical acetyl-lysine (KAc) binding pocket occupied by (+)-JQ1 (**Fig. 1D**). Collectively, the docking results indicate the high binding affinity and selectivity of ZL0580 for BRD4 BD1 at this noncanonical binding site, distinct from the classical KAc binding pocket occupied by JQ1.

**Figure 1.**
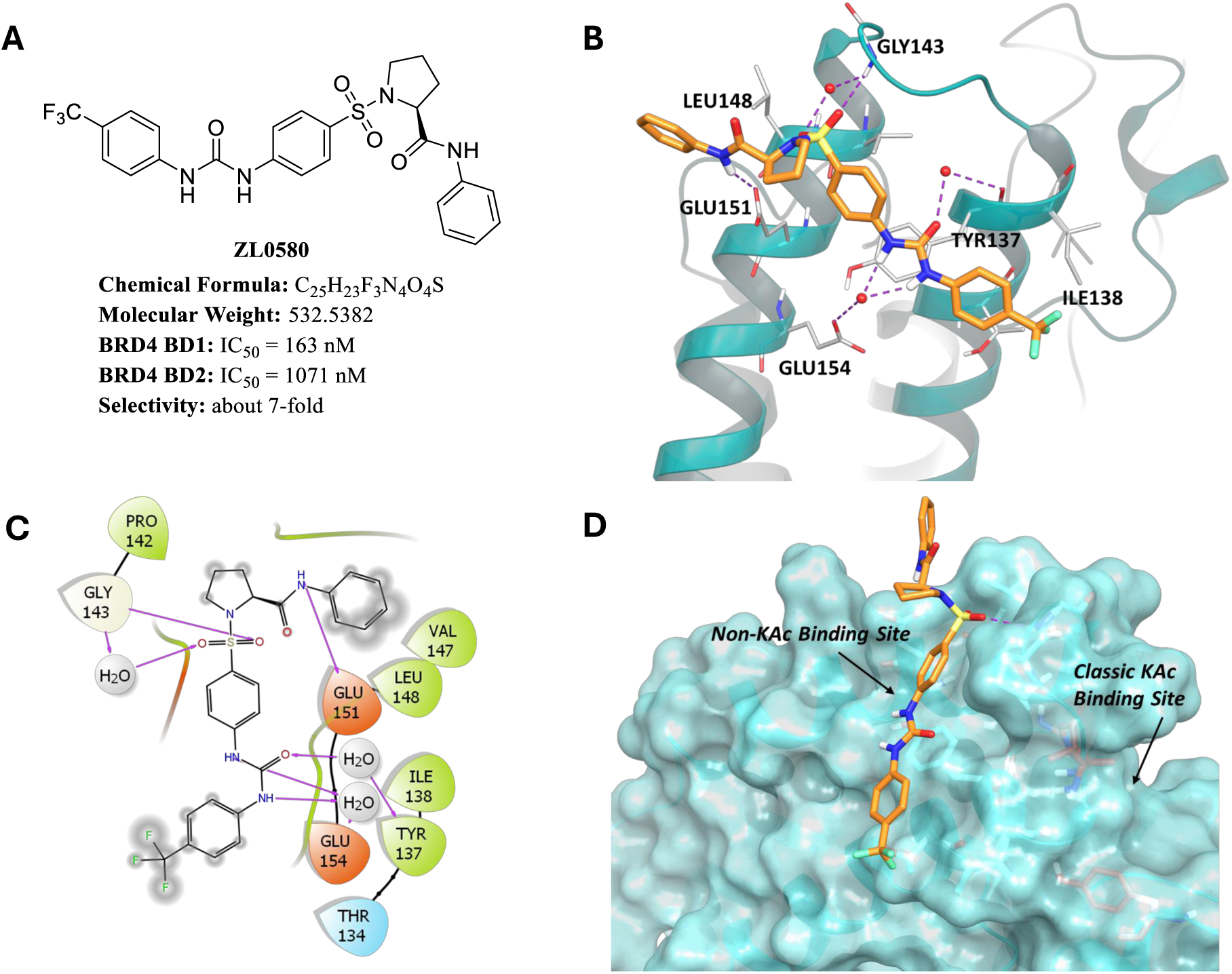
Predicted interactions between ZL0580 and BRD4 BD1. **(A)** Chemical structure of ZL0580 and its in vitro binding affinities toward BRD4 BD1 and BD2 domains. **(B)** Ribbon representation showing ZL0580 (orange stick) docked into the non-KAc binding site of BRD4 BD1 (based on PDB ID: 6U0D), forming strong hydrogen bond interactions (purple dotted lines) with key residues Glu151, Glu154, Tyr137, Gly143, and Leu148. **(C)** 2D interaction diagram illustrating the hydrogen bonding interactions (purple arrows) between ZL0580 and residues at the new binding site of BRD4 BD1. **(D)** Surface representation of BRD4 BD1 showing the binding of ZL0580 at the non-KAc binding site, highlighting its spatial orientation and interaction surface.

### Mutagenesis identifies key residue mediating ZL0580 binding to BRD4 BD1

To confirm that ZL0580 targets a novel binding site distinct from the classical KAc binding pocket of BRD4 BD1 occupied by (+)-JQ1, we performed mutagenesis of key residues located at the new binding site in BRD4 BD1 that was predicted by molecular docking. Docking analysis indicated that GLU151 residue with a polar side chain forms a critical hydrogen bond with the amide NH group ZL0580 (**Fig. 1**). Therefore, GLU151 was selected for mutagenesis, consistent with our previous study demonstrating that ZL0590 binds to a similar noncanonical binding site as revealed by its co-crystal structure with BRD4 BD1 (PDB ID: 6U0D) [42]. The BRD4 BD1 mutant, in which GLU151 was substituted with alanine (E151A), was expressed and purified for thermal shift assay (TSA) to analyze the binding affinity with ZL0580 or JQ1 (**Fig. 2A**).

**Figure 2.**
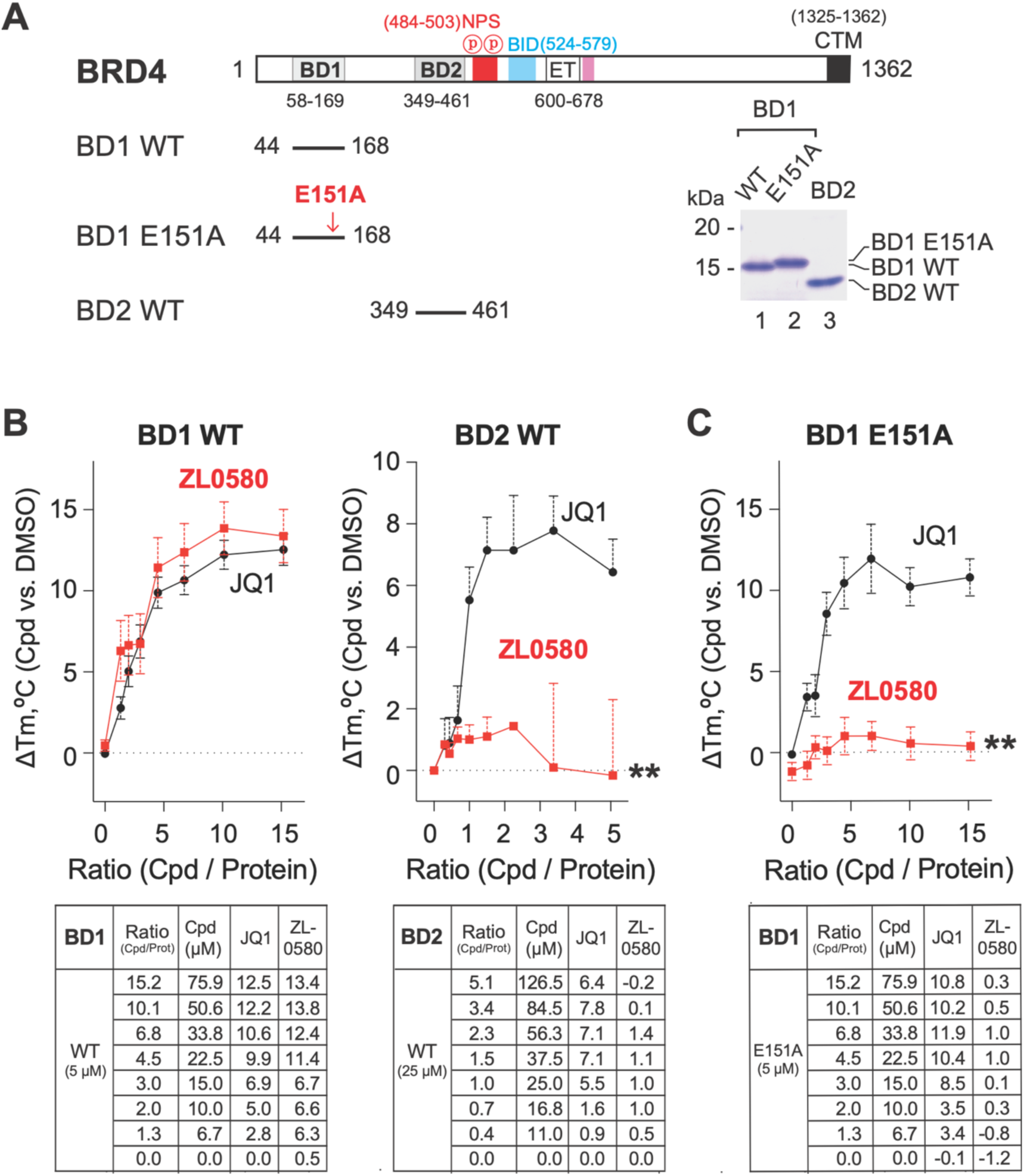
Mutagenesis identifies key residues for selective binding of ZL0580 to BRD4 BD1. **(A)** Schematic representation of BRD4 domain organization of full-length BRD4, indicating the positions of bromodomains (BD1, BD2), N-terminal phosphorylation sites (NPS), basic residue-enriched interaction domain (BID), extra-terminal domain (ET), and C-terminal motif (CTM), with numbers denoting amino acid residues [58]. Constructs used for mutagenesis, including wild-type BD1 (BD1 WT), BD1 with an E151A mutation (BD1 E151A), and wild-type BD2 (BD2 WT), where the red arrow indicates the position of the E151A mutation in BD1. SDS-PAGE analysis showing the expression and purity of recombinant BD1 WT, BD1 E151A, and BD2 WT proteins, with molecular weights (kDa) indicated. **(B)** Thermal stabilization of BRD4 bromodomains by JQ1 and ZL0580 is shown through thermal shift assay (TSA) curves, depicting the change in melting temperature (ΔTm) of BD1 WT (left) and BD2 WT (right) upon titration with increasing concentrations of JQ1 (black circles) and ZL0580 (red squares). Data are presented as mean ± SEM, normalized to DMSO control. **(C)** The effect of E151A mutation on ZL0580 binding to BRD4 BD1 is further illustrated by a TSA curve showing the ΔTm of BD1 E151A upon titration with JQ1 (black circles) and ZL0580 (red squares). Data are presented as mean ± SEM, normalized to DMSO control. ** p<0.01 (ZL0580 compared to JQ1 in B-C).

Both ZL0580 and JQ1 bound to wild-type (WT) BRD4 BD1 (**Fig. 2B**). However, ZL0580 completely lost binding to the BRD4 BD1 E151A mutant compared to WT BRD4 BD1 (**Fig. 2C**), whereas (+)-JQ1 retained comparable binding affinity for the E151A mutant and WT BRD4 BD1 (**Fig. 2B-C**). As an additional control, ZL0580 showed no detectable binding to WT BRD4 BD2 (**Fig. 2B**), further confirming its BD1 selectivity. These results demonstrate that GLU151 is essential for ZL0580 binding and further support that ZL0580 targets a novel binding site on BRD4 BD1, distinct from the classical KAc pocket recognized by (+)-JQ1. This surface-exposed binding site may enable ZL0580 to more efficiently modulate critical protein-protein interactions (PPIs) between BRD4 and its partner proteins [38].

### Opposing transcriptomic profiles induced by ZL0580 and JQ1

Our previous studies showed that although ZL0580 and JQ1 both target BRD4, they exert opposing effects on HIV proviral transcription and latency [38, 41]. To gain further mechanistic insights and assess their global transcriptomic impact, we performed RNA-Seq analysis on latently HIV-infected J-Lat cells treated with ZL0580, JQ1, or with DMSO as the negative control (NC). 24 hours after treatment, RNA was extracted from cells for sequencing analysis. RNA-Seq was conducted on duplicate samples for each condition. Principal component analysis (PCA) demonstrated that ZL0580 induced a distinct but less pronounced transcriptional shift relative to NC compared to JQ1 (**Fig. 3A; Fig. S1)**. Differential gene expression analysis supported this observation (**Fig. 3B**): ZL0580 significantly regulated 767 genes (405 upregulated and 362 downregulated), whereas JQ1 affected 4,378 genes (1,713 upregulated and 2,665 downregulated) using a threshold of p < 0.001 and log2 fold change > 0.5 (**Fig. 3B**). The narrower transcriptomic impact of ZL0580 likely reflects its selective modulation of BRD4, in contrast to JQ1, which targets all four BET family members [43].

**Figure 3.**
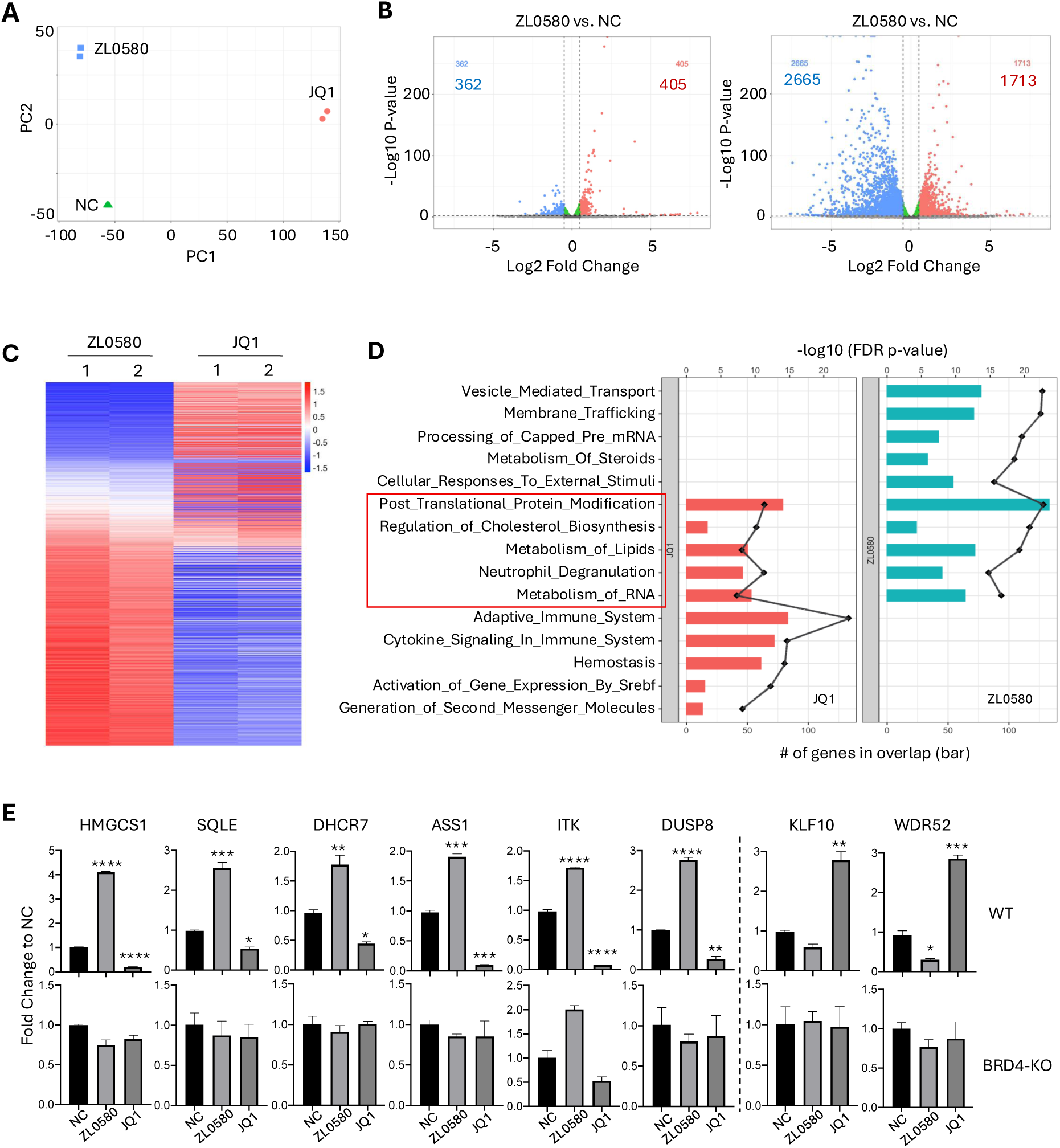
Transcriptomic profiling and pathway analysis of JQ1 and ZL0580 treatments. **(A)** Principal Component Analysis (PCA) plot showing distinct clustering of samples treated with ZL0580, JQ1, and the negative control (NC), indicating differential global gene expression profiles. **(B)** Differential gene expression analysis with volcano plots between ZL0580 vs. NC (left) and JQ1 vs. NC (right). Red dots indicate significantly upregulated genes, blue dots indicate significantly downregulated genes, and green dots represent non-significant genes. The number of upregulated and downregulated genes is shown in red and blue, respectively. **(C)** Heatmap representing the differentially expressed genes and the expression patterns of genes significantly regulated by ZL0580 and JQ1 treatments relative to NC. Red represents upregulation and blue represents downregulation. Samples from each condition are shown duplicates. **(D)** Gene ontology (GO) enrichment analysis shows a bar graph of biological processes significantly enriched among differentially expressed genes in JQ1 (left, red) and ZL0580 (right, blue) treatments. The length of the bars indicates the number of overlapping genes. The black line plot superimposed on each bar graph represents the log10 (FDR-adjusted p-value). Selected pathways with functional relevance (highlighted by a red box) include lipid metabolism, RNA metabolism, cholesterol biosynthesis, and protein modification. **(E)** Quantitative RT-PCR validation of selected differentially expressed genes. Each bar represents the fold change in expression relative to NC for ZL0580 and JQ1 treated samples. Data is shown as mean ± SEM from three replicates. Statistical comparison was performed using one-way ANNOVA. * p<0.05; ** p<0.01; *** p<0.001; **** p<0.0001

Analysis of differentially expressed genes revealed largely inverse transcriptomic profiles induced by ZL0580 and JQ1. Heatmap analysis showed that most genes downregulated by ZL0580 were upregulated by JQ1, and vice versa (**Fig. 3C**), consistent with their opposing effects on HIV transcription and latency [38, 41]. Gene Set Enrichment Analysis (GSEA) showed that both compounds regulate similar biological pathways, with 5 of the top 10 enriched pathways shared between them, including post translational protein modification, regulation of cholesterol biosynthesis, metabolism of lipids, neutrophil degranulation, and RNA metabolism (**Fig. 3D**). Further GSEA analysis indicated that JQ1 also strongly regulated multiple immune-related pathways (**Fig. S2**), whereas ZL0580 had minimal or no effect on these immune pathways **(Fig. S2)**. This observation aligns with our previous finding that ZL0580 minimally affects immune activation and cytokine production in human T cells and PBMCs [38]. Collectively, these results indicate that ZL0580 induces more selective and less extensive transcriptomic changes than JQ1, and that the two compounds trigger largely opposing gene expression programs in J-Lat cells.

To validate the RNA-Seq data, RT-qPCR analysis was performed on a subset of differentially expressed genes identified (**Fig. 3E**). In WT J-Lat cells, ZL0580 upregulated genes such as HMGCS1, SQLE, DHCR7, ASS1, ITK, and DUSP8, whereas these genes were downregulated by JQ1 relative to NC. Conversely, KLF10 and WDR52 were downregulated by ZL0580 but upregulated by JQ1, consistent with the RNA-Seq findings (**Fig. 3E**, top). In the BRD4-knockout (KO) J-Lat cells generated using CRISPR/Cas9 and reported in our previous study [38], the contrasting regulatory effects of ZL0580 and JQ1 on these representative genes were largely abolished, further supporting that BRD4 is a primary target mediating the transcriptomic effects.

### *In vivo* pharmacokinetics and toxicity profile of ZL0580 in mice

Given the favorable *in vitro* target engagement and HIV-suppressive activity of ZL0580, we next evaluated *in vivo* pharmacokinetic (PK) properties and safety profile of the compound in mice prior to efficacy testing. In ICR mice, intravenous administration of ZL0580 (IV, 10 mg/kg) resulted in a favorable PK profile, characterized by high *C*_max_ and AUC values (*C*_max_ = 19259.0 ± 2792.5 ng/mL; AUC_0-last_ = 14428.9 ± 3469.2 ng.h/mL; AUC_0-∞_ = 14442.6 ± 3472.3 ng.h/mL) (**Fig. 4A, C**). Oral administration of ZL0580 (20 mg/kg) showed an acceptable moderate bioavailability (*F* = 38.71 ± 13.03%) (**Fig. 4B, C**).

**Figure 4.**
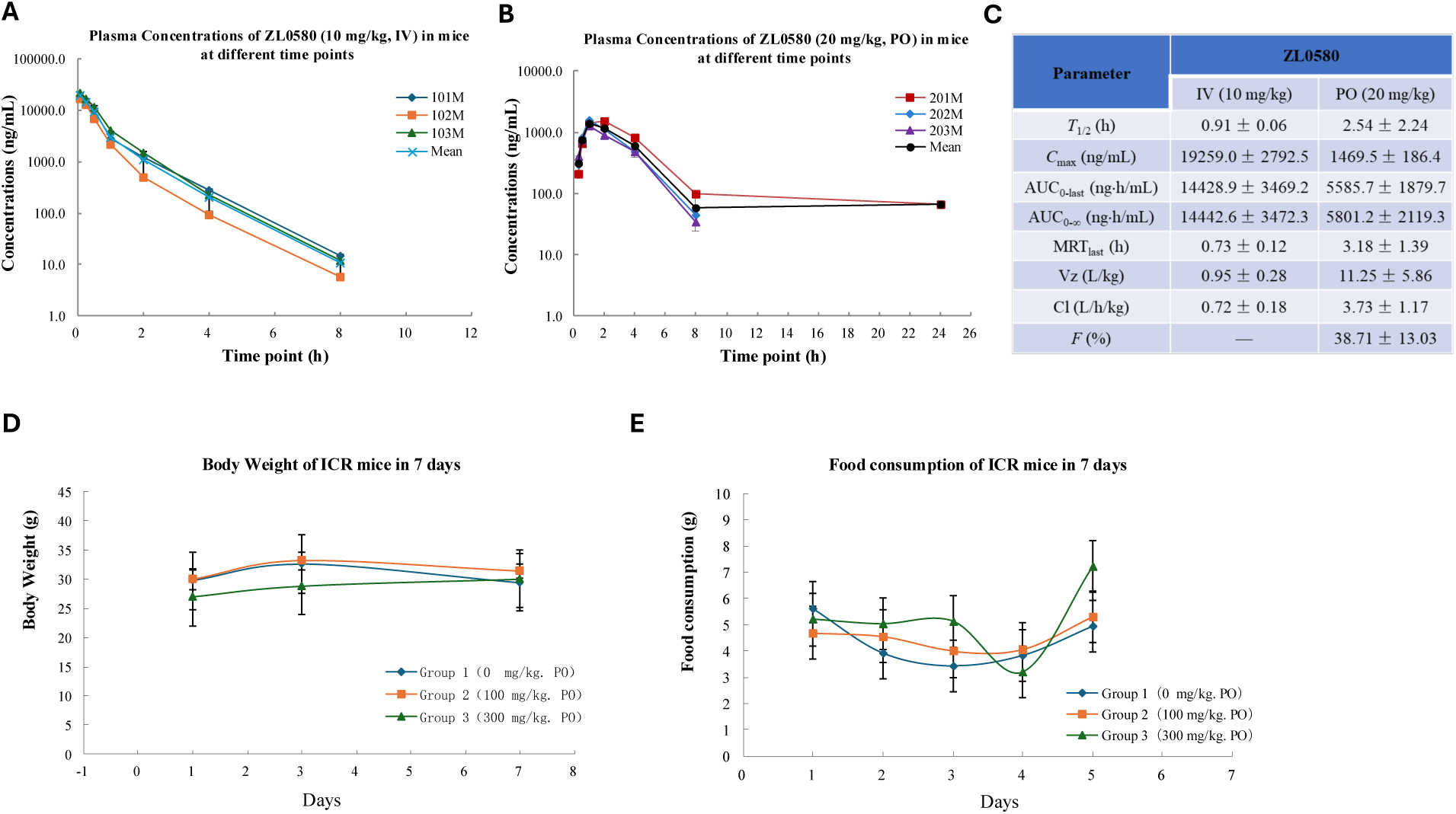
In vivo pharmacokinetics (PK) and toxicity profiles of ZL0580 in ICR mice. **(A)** Plasma concentrations of ZL0580 following intravenous administration at 10 mg/kg in male ICR mice (n=3 per group) measured at various time points. **(B)** Plasma concentrations of ZL0580 following oral administration at 20 mg/kg in male ICR mice (n=3) over time. **(C)** Summary of in vivo PK parameters of ZL0580 in male ICR mice. **(D)** Body weight changes in ICR mice over the treatment period (n=10 per group). (E) Food consumption changes in ICR mice monitored throughout the study (n=10 per group).

Toxicity assessments in mice revealed that ZL0580 was well tolerated *in vivo* (**Fig. 4D-E**). Daily administration at doses of 0 mg/kg, 100 mg/kg, or 300 mg/kg for 7 days caused little to no changes in body weights (**Fig. 4D**) or food consumption (**Fig. 4E**). No clinical signs of toxicity were observed throughout the study (**Table S1**). Serum biochemistry as well as necropsy and gross pathological examinations revealed no abnormalities at the end of the study (**Table S2-S3**). Collectively, these findings demonstrate that ZL0580 has favorable PK properties and an acceptable safety profile in mice, supporting its further evaluation in HIV-suppression studies *in vivo*.

### *In vivo* activity of ZL0580 to suppressing HIV in HIV-infected humanized mice

To assess *in vivo* HIV-suppressive activity of ZL0580, we utilized a humanized mouse model (Hu-mice) generated by engrafting human CD34+ hematopoietic stem cells (HSCs) into the immunodeficient NOD SCID gamma (NSG) mice (the Jackson Laboratory). Twelve Hu-mice, aged 16–20 weeks and engrafted with human HSCs from a single donor, were enrolled in the study. Prior to HIV infection, human immune cell reconstitution was evaluated in all animals. Human CD45+ cells were detected in the peripheral blood of all 12 Hu-mice, ranging from 36% to 62% with a median of 45% **(Fig. S3A)**. Further characterization revealed successful reconstitution of major human immune cell subsets **(Fig. S3B)**. Within the human CD45+ population, the median proportions of CD3+ T cells, CD19+ B cells, and CD33+ myeloid cells were 14.9%, 74.3%, and 3.7%, respectively, confirming robust human hematopoietic reconstitution (**Fig. S3B**).

Next, at week 0, all12 Hu-mice were intravenously infected with a single dose of HIV-1 JR-CSF (equivalent to 10 ng HIV p24 antigen per animal) (**Fig. 5A**). Systemic HIV infection was confirmed in 11 out of the 12 mice at weeks 2 and 3, based on the detection of plasma viral RNA quantified by a single-step RT-qPCR (**Fig. 5B-C**). Systemic HIV infection in Hu-mice was also confirmed by ELISA quantification of plasma HIV p24 antigens at weeks 2 and 3 after HIV inoculation (**Fig. S4**). The 11 infected Hu-mice were then randomized into four treatment groups: empty vehicle control (EV; n=2), ART (n=3), ZL0580 (n=3), and ART+ZL0580 (n=3). Drug formulations included: ART (Raltegravir 1.2 mg/mouse, Emtricitabine 3.2 mg/mouse, Tenofovir 3.2 mg/mouse), ZL0580 (3.8 mg/mouse), and a combination of both ART+ZL0580. All drugs were formulated in the buffer consisting of 10% DMSO, 10% Solutol HS 15, and 80% HP-β-CD (20%, w/v). The empty vehicle (EV) group received only the formulation buffer. Treatment regimens were initiated at week 3 and administered daily by intraperitoneal injection (**Fig. 5A**). Plasma viral loads were closely monitored during the treatments.

**Figure 5.**
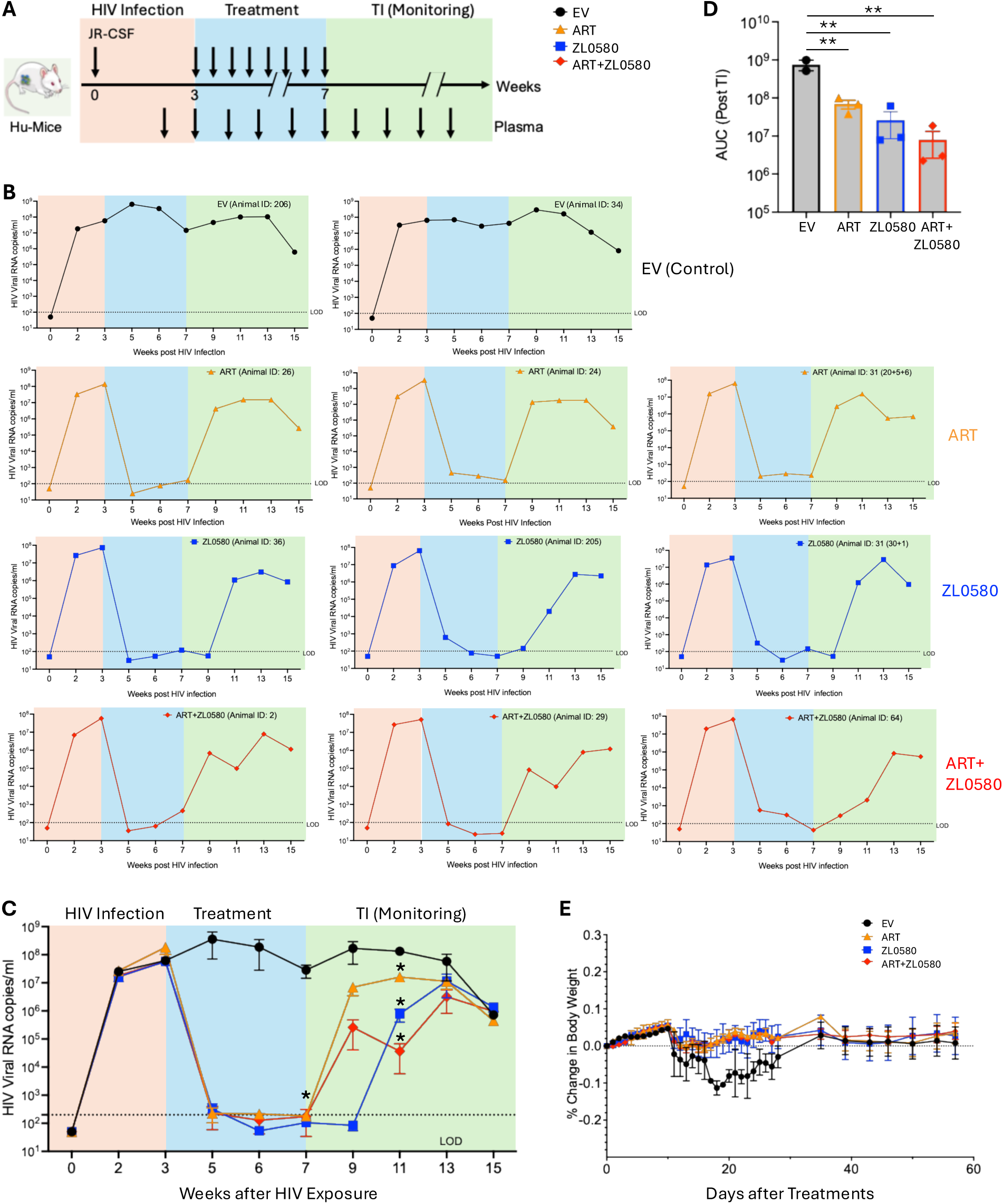
In vivo activities of treatments to suppress HIV in humanized mice. **(A)** Experimental design and timeline for the *in vivo* study using HIV-infected CD34+ hu-mice. The timeline shows when HIV infection occurred, the duration of treatment, and the period of treatment interruption (TI) with subsequent monitoring. Treatment groups included Empty Vehicle (EV; n=2), Antiretroviral Therapy (ART; n=3), ZL0580 (n=3), and ART + ZL0580 (n=3). Blood/plasma samples were collected as indicated for the quantification of HIV viral loads. **(B)** Kinetics of plasma viral loads (HIV RNA copies/ml) post HIV infection for each mouse in the Empty Vehicle (top row), ART (second row), ZL0580 (third row), and ART + ZL0580 (bottom row) treatment groups. The shaded areas denote the phase of establishing systemic HIV infection (orange), treatment phase (blue), and treatment interruption (TI) monitoring phase (green). RT-qPCR was performed in duplicate and the mean for each sample at different time points was shown. **(C)** Summary data for kinetics of plasma viral loads over weeks post HIV Infection for all animals in the four treatment groups: Empty Vehicle (black circles), ART (orange triangles), ZL0580 (blue squares), and ART + ZL0580 (red diamonds). Data are presented as mean ± SEM. Statistical comparison of viral loads among the four groups at each time point was performed using one-way ANNOVA. * at week 7 and *** at week 13 denote significant difference of the individual treatment group (ART, ZL0580, or ART+ZL0580) as compared to the EV group. The dashed line indicates the limit of detection (LOD). **(D)** Comparison of area under the curve (AUC) among the four groups after treatment interruption (AI) between week 7 and 15. Statistical comparison was performed using one-way ANNOVA. **(E)** Percentage change in body weight over days post treatment (DPT) for all treatment groupsData are presented as mean ± SEM. In this Figure: * p<0.05; ** p<0.01

The results showed that in the EV control group, plasma viral loads were consistently high throughout the study in both animals (**Fig. 5B-C**). In contrast, all three treatment groups (ART alone, ZL0580 alone, or ART+ZL0580) demonstrated drastic reductions in plasma viremia. By week 5 (after two weeks of daily treatment), the plasma viral loads were reduced to nearly undetectable levels in all mice of all the three groups, including the ZL0580 monotherapy group (**Fig. 5B**), providing strong evidence that ZL0580 alone as a host-directed compound robustly suppresses active HIV replication *in vivo*. The low-to-undetectable HIV viremia in persisted through weeks 7, during which all mice remained under treatments in the three groups (**Fig. 5B**). At week 7, the differences in plasma viral loads for the three treated groups (ART, ZL0580, ART+ZL0580) compared to the EV control group reached statistical significance (p<0.05) (**Fig. 5C**).

To assess viral rebound following treatment cessation, an analytical treatment interruption (TI) was initiated at week 7 across all the groups (**Fig. 5A**). Plasma viral loads were subsequently measured once every two weeks from week 7. At week 9 (two weeks post-TI), rapid viral rebounds were observed in all three Hu-mice in the ART group (mean viral copies: 6.96 × 10^6^ copies/mL), despite the levels remained much lower than the EV group (mean viral copies in EV: 1.75 × 10⁸ copies/mL) (**Fig. 5C**). Notably, all mice in the ZL0580 alone group maintained undetectable viremia at week 9 (**Fig. 5C**), suggesting that compared to ART, ZL0580 could further delay viral rebounds. At week 11 (four weeks post-TI), plasma viremia became detectable in all mice in both ZL0580 and ART+ZL0580 groups; however, viral loads in these two groups at this time point remained markedly lower than those in the ART group: mean viral RNA copies of 7.7 × 10⁵ copies/mL in ZL0580 group and of 3.6 × 10⁴ copies/mL in ART+ZL0580 group, as compared to mean viral RNA copies of 1.6 × 10⁷ copies/mL in ART group. Statistical analysis showed that, at week 11, the plasma viral loads in all three treated groups were significantly lower than that in the EV group (**Fig. 5C**). By week 15 (eight weeks post-ATI), viral loads in all the treatment groups converged to similar levels (**Fig. 5C**). To more quantitatively assess the impact of different treatments on viral rebounds after TI, we quantified area under the curve (AUC) between weeks 7 and 15 as shown in **Fig. 5C**, which combines the parameters of time (x-axis) and viral loads (y-axis) after TI. The data demonstrated that the three treatments (ART, ZL0580, and ART+ZL0580) induced significant effects on repressing viral rebounds compared to the EV control group (**Fig. 5D**).

To assess general health of the mice and potential protective effects of the treatments, body weights of Hu-mice were closely monitored. As shown in **Fig. 5E**, all treatment groups, including those receiving ART, ZL0580, or combined ART+ZL0580, maintained stable body weights with no significant deviations throughout the study. There was a slight reduction in body weights of Hu-mice in the EV group following HIV infection. However, no statistically significant difference was detected as compared to the three treatment groups (**Fig. 5D**). Nonetheless, these data demonstrate that daily intraperitoneal administration of ZL0580 alone, or in combination with ART, is well tolerated in Hu-mice over the course of treatment. These results provide *in vivo* evidence that ZL0580 robustly suppresses active HIV replication, reducing plasma viral loads to nearly undetectable levels, and modestly delays viral rebound following treatment interruption.

### HIV DNA in blood cells across different groups

To assess whether ZL0580 or ART affects HIV DNA levels, we quantified cell-associated HIV DNA in the peripheral blood of Hu-mice after treatments. Blood cells collected at weeks 7 and 13 were subjected to DNA extraction and quantification of HIV DNA by qPCR. The results showed that no significant difference in cell-associated HIV DNA copies was detected among the EV control, ART, ZL0580, and the ART+ZL0580 groups at week 7 (**Fig. 6A**) or week 13 (**Fig. 6B**). The data indicate that while ZL0580 and ART robustly suppress active viral replication (as reflected by plasma viral RNA levels), neither treatment significantly alters the levels of HIV DNA in peripheral blood of Hu-mice.

**Figure 6.**
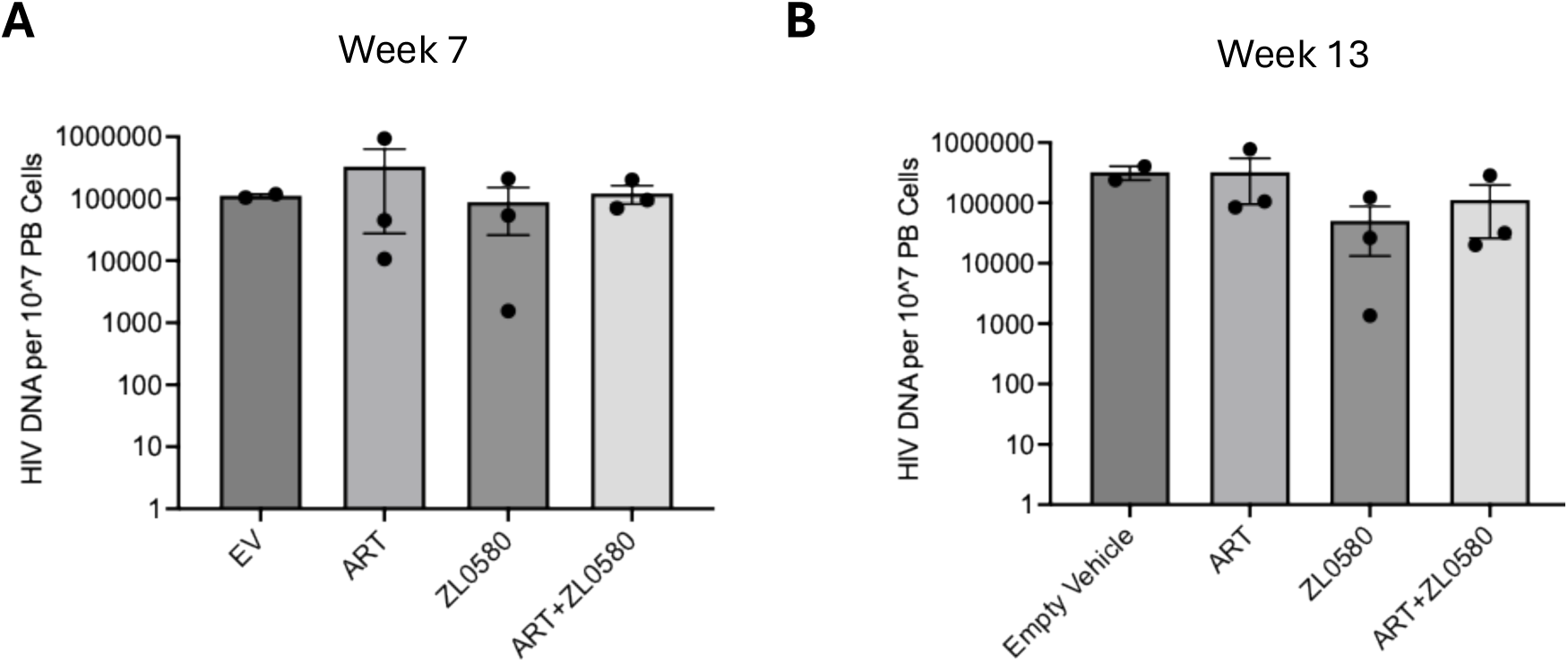
HIV DNA copies in blood cells. HIV DNA copies per 10^7^ peripheral blood (PB) cells at week 7 (A) and week 13 (B) for Empty Vehicle (EV), Antiretroviral Therapy (ART), ZL0580, and ART+ZL0580 treatment groups. Data are presented as mean ± SEM, with individual data points shown.

## DISCUSSION

The “block and lock” strategy represents a promising approach for achieving a functional HIV cure [18–20]. In this study, we evaluated bioavailability and *in vivo* HIV-suppressive activity of the BRD4-targeting small molecule ZL0580 in animal models. ZL0580 was bioavailable and exhibited a favorable safety profile, with no evident toxicity in mice. In a humanized mouse model, ZL0580 monotherapy potently suppressed active HIV replication and modestly delayed viral rebound after treatment cessation. To our knowledge, this is the first demonstration of robust *in vivo* HIV suppression by a small-molecule compound targeting host epigenetic machinery. Further, we identified key residues mediating the selective interaction of ZL0580 with BRD4 BD1, providing new mechanistic insights into its distinct modulation of HIV transcription and latency compared with the pan-BET inhibitor JQ1.

Our previous studies demonstrated that ZL0580 and JQ1 exhibit distinct binding profiles within the BET protein family, with ZL0580 selectively targeting BRD4 BD1, whereas JQ1 non-selectively binds both BD1 and BD2 of all BET proteins [38, 40, 41]. The molecular basis underlying this selectivity was unclear. The present study identified the glutamic acid 151 (E151) in BRD4 BD1 as a key residue mediating selective ZL0580-BRD4 interaction. This finding corroborates previous structural evidence supporting BRD4 BD1 as a druggable site [44] and confirms that ZL0580 engages a distinct binding interface, avoiding the conserved acetyl-lysine (KAc) binding pocket targeted by pan-BET inhibitors such as (+)-JQ1 [38, 41, 45]. The selective targeting of BD1 by ZL0580, and its preferential engagement of BRD4 over other BET proteins, confers a pharmacological profile that may minimize disruption of global transcriptional networks, a limitation of the first-generation BET inhibitors [41, 42, 45].

RNA-seq analysis of ZL0580-and JQ1-treated cells revealed several interesting results. First, modulation of BRD4 by ZL0580 or JQ1 results in both upregulation and downregulation of gene expression. This is consistent with previous studies reporting that BRD4 inhibition - via either genetic knockout [46] or pharmacologic inhibition [47] - can activate or repress transcription in a gene-specific manner, highlighting the dual role of BRD4 as a transcriptional activator [48] or repressor [49]. Second, while JQ1 modulated the expression of a broad array of cellular genes (>4000), ZL0580 elicited a more selective and attenuated transcriptional response, affecting approximately 760 genes. This is likely attributable to the selective targeting of BRD4 by ZL0580, in contrast to JQ1’s pan-BET inhibition profile [43]. The restrained transcriptional footprint of ZL0580 supports a notion that selective BD1 inhibition may allow for more targeted repression of viral transcription while sparing host gene regulation critical for homeostasis [50]. Third, gene enrichment analysis reveals that multiple pathways were commonly regulated by both compounds but the gene expression within these pathways were inversely regulated. This aligns with their opposing effects on HIV transcription [38, 40, 41] and reinforces the idea that that both compounds modulate overlapping molecular targets through distinct mechanisms. Lastly, ZL0580 induced minimal to changes in immune and pro-inflammatory gene expression pathways, corroborating previous finding that ZL0580 does not perturb immune activation and cytokine production in human T cells and PBMCs [38]. This contrasts with the widespread transcriptional regulation of immune-related pathways by JQ1, which has been linked to cytokine release and immunomodulatory effects [51, 52].

In the humanized mouse model of HIV infection, ZL0580 monotherapy robustly suppressed active HIV replication and reduced plasma viremia to nearly undetectable levels after two weeks of daily treatment in Hu-mice. This *in vivo* result is highly consistent with the *in vitro* and *ex vivo* evidence reported in our previous studies using cell line and primary cell models [38, 40, 41]. Mechanistically, by inhibiting Tat-mediated recruitment of positive transcription elongation factor b (p-TEFb) and preventing transcriptional elongation, ZL0580 induces a transcriptionally inert state at the HIV long terminal repeat (LTR) [38]. This HIV-suppressive mechanism of ZL0580 is further reinforced by its ability to stabilize a repressive chromatin structure at the proviral promoter [38]. In this Hu-mouse model of HIV infection, ZL0580 treatment also delayed viral rebound following TI, compared to ART alone, despite the effect was modest (∼2 weeks). Of note, after viral rebounds, the plasma viral loads in the ZL0580 and ZL0580+ART groups remained markedly lower than those in the ART group. These findings together provide early *in vivo* proof-of-concept evidence for ZL0580 as a potential HIV “block and lock” candidate. However, the lack of durable viral suppression by ZL0580 after treatment cessation highlights a limitation of the compound, which is likely attributed to its relatively short half-life. Future research should be conducted to enhance the half-life and durability of HIV suppression by this class of molecules. Additionally, combining multiple “block and lock” agents with distinct modes of action is also a promising approach to achieve more durable HIV suppression after treatment cessation.

Indeed, the latency-promoting potential of ZL0580 is enhanced when used in combination with LEDGINs, as demonstrated by Pellaers et al. [23]. LEDGINs function by retargeting HIV provirus into transcriptionally inactive genomic regions, thereby minimizing the likelihood of proviral reactivation [21, 22]. The synergistic effects between ZL0580 and LEDGINs support a potential combinatorial “block-and-lock” strategy aimed at enforcing deep and durable latency [23]. This dual-targeting approach addresses both integration site bias and the maintenance of transcriptional repression of the integrated provirus. Previous studies have shown that LEDGINs promote integration into genomic regions less permissive to transcriptional reactivation [22, 53]. Nevertheless, residual low-level expression may persist from certain integration sites, highlighting the need to combine LEDGINs with epigenetic silencers such as ZL0580. The complementary mechanisms of these latency-promoting agents (LPAs) highlight the importance of concurrently targeting both the chromatin context and transcriptional machinery to achieve durable HIV silencing.

Given the duration and complexity of humanized mouse studies involving chronic HIV infection and continuous drug administrations, a limitation of the present study is the relatively small number of animals used. Nonetheless, the low inter-animal variability within each group enabled the detection of statistically significant differences in plasma viremia among groups in our study. These proof-of-concept findings warrant further confirmation and validation in larger-scale animal studies and non-human primate (NHP) models.

In summary, our study supports that ZL0580 is a BD1-selective BRD4 modulator with mechanistic and functional distinctions from the classical pan-BET inhibitors. Its ability to suppress HIV transcription, while minimizing global disruption of host chromatin architecture and immune activation, positions it as a valuable tool and therapeutic candidate for HIV latency-promoting strategies. Future efforts should be pursued to improve the half-life and durability of HIV suppression by this class of molecules. Additionally, combinatorial approaches incorporating multiple LPAs with distinct mechanisms—such as epigenetic silencers (ZL0580), LEDGINs, and Tat inhibitors — are likely necessary to achieve maximal and sustained HIV suppression or silencing.

## MATERIALS AND METHODS

### Molecular Docking Studies

The molecular docking study was performed using the Schrödinger Small-Molecule Drug Discovery Suite. The crystal structure of compound ZL0590 in complex with BRD4 BD1 (PDB ID: 6U0D) was downloaded from RCSB PDB Bank and prepared with Protein Prepared Wizard. During protein preparation, hydrogens were added, crystal waters were removed while water molecules around the KAc pocket were maintained, and other parameters were used by default. The 3D structure of ZL0580 was generated with Schrödinger Maestro, and the initial lowest energy conformation was calculated with LigPrep. For the docking study, the grid center was chosen on the centroid of the KAc site occupied by ZL0590 in the cocrystal structure and a 24 × 24 × 24 Å grid box size was fixed. The docking was performed with Glide using the XP protocol. The docking pose was incorporated into Schrödinger Maestro for the ligand-receptor interaction analysis.

### Protein Expression and Purification

Plasmid construction and subsequent protein expression and purification of BRD4 proteins were performed using OverExpress C43(DE3) bacterial competent cells (Lucigen) as described in our previous studies [42, 54]. In brief, 12 L of bacterial cultures grown to OD600 between 0.6-0.8 were induced at 37°C for 4 h by 1 mM isopropyl β-bd-1-thiogalactopyranoside (IPTG). Bacterial sonication supernatant in the lysis buffer (500 mM NaCl, 20 mM Tris-HCl pH 7.9 at 4°C, 20% glycerol, 1 mM EDTA, 0.2 mM DTT, and protease inhibitors) was incubated with glutathione (GSH) agarose beads (GOLDBIO, G-250-100) overnight with rotation at 4°C. The protein-bound GSH beads were washed three times with the lysis buffer and digested with human rhinovirus 3C protease (HRV-P3C; Thermal Fisher 88946) at 4°C for 4 h, according to the manufacture’s procedure, to remove GST tag from BD1/2 proteins. Untagged BRD4-BD1 and -BD2 were separated from HRV-P3C via passing through a HiTrap Q HP anion exchange chromatography column (GE Healthcare, GE17-1154-01) and eluted by a salt gradient from 100 mM to 1 M NaCl. The peak fractions were combined and concentrated using Amicon Ultra Centrifugal Filter with a molecular weight cut-off of 3 kDa (Millipore, UFC900324) in the final buffer 10 mM HEPES pH 7.5 and 150 mM NaCl.

### Thermal Shift Assay (TSA)

TSA experiments were performed in triplicate in a 384-well plate (Bio-Rad, HSP3801) using the CFX384 Real-Time C1000 touch thermal cycler (Bio-Rad) as described previously [42, 54]. In each 10-μL reaction, 5 μM of WT and mutant BD1 and 25 μM of BD2 was mixed with 2 μL of 25X Sypro Orange protein dye (Invitrogen, S6650) in 20 mM HEPES pH 6.5, 100 mM NaCl, and 2.5% glycerol with 2 μL of ZL580 or JQ1 in 5% DMSO (final 0–126.5 μM). Fluorescence signals generated by dye binding to unfolded proteins were monitored by using the melt curve protocol at the rate of 1°C temperature increment in 30 s from 25 °C to 95 °C with signal capture every 1 °C. The melt curves obtained by fluorescence signal (F) versus temperature (T) were converted to melting peaks using Bio-Rad CFX Manager v3.1 software. Tm is the temperature at which 50% of protein is unfolded and bound by the dye. ΔTm is the difference of Tm comparing compound-treated vs. untreated (i.e., DMSO-only) samples.

### Pharmacokinetics (PK) studies

Animal experiments assessing PK and *in vivo* toxicity profile of ZL0580 was performed by the contracting research organization (CRO) Sundia. The studies were performed according to the NIH Guide for Care and Use of Experimental Animals. The PK studies of ZL0580 was performed using male ICR mice (18-22 g). The mice were randomly divided into two groups (*n*=3). The mice of the group 1 were intravenously administrated with ZL0580 [10 mg/kg, in 10% DMSO + 10% Solutol HS 15 + 80% HP-β-CD (20%, w/v)] and the mice of the group 2 were orally administrated with ZL0580 [20 mg/kg, in 10% DMSO + 90% HP-β-CD (20%, w/v)]. Blood samples of group 1 and group 2 mice were collected at the indicated time points post treatment and put into heparinized centrifugation tubes, which were centrifuged under the condition of 6800 rpm and 4 °C for 6 min. The supernatant was collected and stored at -20 °C for further LC-MS analysis. Before the plasma concentrations (AUC) were determined, standard curves were generated with various concentrations of ZL0580 and the internal standard (IS). All PK parameters were calculated with WinNonlin® 6.4 software.

### *In vivo* toxicity studies

The acute toxicity studies were performed in both male and female ICR mice. The mice were divided into 3 groups (each group, n = 10, 5 males and 5 females). Group 1 was the vehicle group without ZL0580 treatment (po, 20% DMA + 20% Solutol HS15 + 60% HP-β-CD (20%, w/v)). Group 2, and group 3 mice were orally administrated with ZL0580 at doses of 100 mg/kg and 300 mg/kg [in 20% Dimethyl Acetamide (DMA) + 20% Solutol HS15 + 60% HP-β-CD (20%, w/v)], respectively. The body weights of all mice were weighed on day 1, 3, and 7 after treatments. The mean body weights of all mice in each group were calculated and the standard curves were generated with the mean body weights of each group mice at different recorded time points. The food consumption masses of each group mice were recorded on day 1, 2, 3, 4, and 5 after treatments. The mean food consumption of all mice in each group were calculated and the standard curves were generated with the mean food consumption of each group at different recorded time points.

### RNA-sequencing

J-Lat cells were treated with 5 µM of either JQ1 or ZL0580 for 24 hours to assess the impact of these compounds on global gene expression. Following the treatment, total RNA was extracted from the cells using the RNeasy Mini Kit (Qiagen) in accordance with the manufacturer’s instructions. To ensure the integrity and quality of the isolated RNA, samples were first quantified and then analyzed by agarose gel electrophoresis. Clear and intact ribosomal RNA bands confirmed that the RNA was of high quality and suitable for downstream applications. The verified RNA samples were subsequently submitted for high-throughput RNA sequencing (RNA-seq) to evaluate transcriptomic changes induced by the respective treatments. RNA sample quality was assessed using Agilent Bioanalyzer and samples with a RIN value larger than 7.0 were used for library preparation. RNA-Seq libraries were prepared using NEBNext Poly(A) mRNA Magnetic Isolation module (E7490) and NEBNext Ultra II Directional RNA Library Prep kit for illumina (E7760) following manufacturer’s recommended procedure. The library quality was assessed using Agilent Bioanalyzer High Sensitivity DNA chip and real-time PCR and pooled together for sequencing. The pooled library was sequenced on Illumina NextSeq 550 Mid Output 150 cycle kit for Paired End 75 bp reads targeting ∼20 million paired reads per sample. The sequencing reads were mapped to human reference genome hg38 using STAR v2.7.5c [55] using recommended ENCODE parameters. Differential gene expression analysis was performed using Bioconductor DESeq 2 package [56].

### Compound formulation for mouse administration

All humanized mice used in this study weighed approximately 20 grams, and drug dosages were calculated accordingly. The following compounds were administered to mice daily: Raltegravir (Ral) at 60mg/kg (1.2 mg/mouse), Emtricitabine (Emt) at 160mg/kg (3.2 mg/mouse), Tenofovir (Ten) at 160mg/kg (3.2 mg/mouse), and ZL0580 at 160mg/kg (3.2 mg/mouse). ART drugs were purchased from MedChemExpress. ZL0580 was synthesized in-house as previously reported [38]. All compounds were dissolved in a formulation vehicle composed of 10% DMSO, 10% Solutol HS 15, and 80% hydroxypropyl-β-cyclodextrin (HP-β-CD, 20% w/v). Compounds were administered via intraperitoneal injection daily.

### Humanized mouse study

#### Ethics Statement

The use of human CD34+ engrafted NSG mice (CD34+ Hu-mice) was approved by the Institutional Animal Care & Use Committee (IACUC) (No.: 2105034) at the University of Texas Medical Branch (UTMB). The fully reconstituted CD34+ Hu-NSG mice were directly obtained from the Jackson Laboratory (Stock No: 005557). The human cells used in the study were de-identified and the study is considered as non-human subject research.

Mice were 16-week-old at the initiation of the experiments. All mice received human CD34+ stem cells of a single donor and human immune cell reconstitution was confirmed for all mice. A total of 12 Hu-mice were initially included in the study with each tagged and assigned with a unique ID. At week 0, all Hu-mice were intravenously (IV) infected with HIV-1 JR-CSF at a single dose of 10ng HIV p24 equivalent virus per animal. Blood/plasma viral loads were monitored and quantified weekly by a single-step RT-qPCR until systemic HIV infection was established. Following HIV exposure, one animal failed to develop systemic infection and was excluded from the cohort. At week 3, the remaining 11 Hu-mice were randomized into four groups for subsequent treatments: Empty Vehicle (EV) (n=2), ART (n=3), ZL0580 (n=3), and ART+ZL0580 (n=3). The control mice received a mixture of DMSO, Solutol HS 15, and hydroxypropyl-β-cyclodextrin (HP-β-CD, 20% w/v), which was considered the empty vehicle control (EV). ART and ZL0580 compounds were administered to mice via intraperitoneal injection daily for four weeks (from week 3 to week 7). Compound dosages were described above. Plasma samples were collected from mice either weekly or biweekly for quantification of viral loads (RNA copies). Blood cells were also collected at weeks 7 and 13 for quantification of cell-associated HIV DNA in blood. Mouse weights were monitored after treatment.

#### HIV p24 ELISA

HIV p24 antigen levels in plasma from humanized mice were quantified using the HIV-1 p24 Antigen Capture Assay Kit (ZeptoMetrix Corporation, USA) according to the manufacturer’s instructions. Briefly, standards and appropriately diluted plasma samples were added to microplate wells pre-coated with anti-p24 antibodies and incubated at room temperature for 2 hours. After washing six times with wash buffer, 100 µL of detector antibody was added and incubated for 1 hour, followed by another six washes. Substrate solution (100 µL) was then added and incubated for 30 minutes at room temperature. Upon color development, 100 µL of stop solution was added, and absorbance was measured at 450 nm using a microplate reader.

#### Plasma viral RNA quantification

Viral RNA was isolated from plasma samples using the QIAamp® Viral RNA Mini Kit (Qiagen) according to the manufacturer’s instructions. Quantification of viral RNA was performed using the PrimeTime™ One-Step RT-qPCR Master Mix (IDT) on a CFX Connect Real-Time PCR System (Bio-Rad). Following RNA extraction, one-step RT-qPCR was carried out using LTR Gag-specific primers and a HEX-labeled probe. Forward: 5′-GATCTCTCGACGCAGGACTC-3′, Reverse: 5′-CGCTTAATACCGACGCTCTC-3′, Probe: 5’-/5HEX/CCAGTCGCC/ZEN/GCCCCTCGC CTC/3IABkFQ. Each 20 µL reaction mixture contained 10 µL of PrimeTime One-Step Master Mix, 1 µL of each primer, 1 µL of probe, and 2 µL of extracted RNA. The thermal cycling conditions were as follows: cDNA synthesis at 50 °C for 15 minutes, polymerase activation at 95 °C for 15 minutes, followed by 44 cycles of denaturation at 95 °C for 15 seconds and annealing/extension at 60 °C for 1 minute. All reactions were performed in duplicate. Following PCR amplification, Viral RNA copies were quantified using the established HIV Gag standard curve as reported in our previous studies [38, 41].

#### Quantification of HIV DNA in blood cells

Quantification of cell-associated HIV DNA in blood cells was performed by qPCR as previously described with modifications [57]. Blood cells were counted and subjected to DNA extraction using the Qiagen QIAamp DNA Blood Mini Kit (QIAGEN). qPCR primers and probe: Forward: 5′-GATCTCTCGACGCAGGACTC-3′, Reverse: 5′-CGCTTAATACCGACGCTCTC-3′, Probe: 5’-/5HEX/CCAGTCGCC/ZEN/GCC CCTCGCCTC/3IABkFQ. The thermal cycling conditions were as follows: 95 °C for 15 minutes, followed by 44 cycles of denaturation at 95 °C for 15 seconds and annealing/extension at 60 °C for 1 minute. All reactions were performed in duplicate. HIV DNA copies were quantified using the HIV Gag standard curve as described above, as well as in a previous study [38], and normalized to peripheral blood cell numbers.

#### Statistics

Statistical analysis was performed using the GraphPad Prism. Data are presented as mean ± SEM where appropriate. Statistical comparison was performed using a non-paired Student’s t test for two groups and a one-way ANOVA for more than two groups. Two-tailed p values were denoted, and p values of less than 0.05 were considered significant.

## ACKNOWLEDGEMENT

The study was funded by NIH grants AI145666 and AI157852 (to H.H). H.H. is supported by NIH grants AI157852 and AI181134. C.-M.C. is supported by NIH grants R01CA288743 and R01CA251698. J.Z. is partly supported by the John D. Stobo, M.D. Distinguished Chair Endowment, and the Edith & Robert Zinn Chair Endowment in Drug Discovery. N.C.H. is supported by a NIH T32 training grant AI060549. The reagents J-Lat cells (clone 10.6, ARP-9849) and JR-CSF HIV molecular clone (ARP-2708) were obtained from the HIV Reagent Program. The content of this paper is solely the responsibility of the authors and does not necessarily represent the official views of the NIH.

## Supporting Information Captions

### Supplementary Figures

Figure S1. Sample-to-sample distance heatmap of gene expression profiles. Figure S2. Pathways regulated by ZL0580 and IQ1.

Figure S3. Human immune cell reconstitution in humanized mice.

Figure S4. Serum HIV p24 concentrations in the individual humanized mice at weeks 2 and weeks 3 after HIV infection.

### Supplementary Tables

Table S1. The result of clinical observation after oral dose of ZL0580 at 100, 300 mg/kg dose levels.

Table S2. Serum Biochemistry in mice after ZL0580 administration.

Table S3. The result of necropsy macroscopic after oral dose of ZL0580 at 100, 300 mg/kg dose levels.

